# Gene annotation bias impedes biomedical research

**DOI:** 10.1101/133108

**Authors:** Winston A. Haynes, Aurelie Tomczak, Purvesh Khatri

**Affiliations:** Stanford Institute for Immunity, Transplantation, and Infection, Stanford University, Stanford, California, USA; Stanford Center for Biomedical Informatics Research, Department of Medicine, Stanford University, Stanford, California, USA; Biomedical Informatics Training Program, Stanford University, Stanford, California, USA

## Abstract

We found tremendous inequality across gene and protein annotation resources. We observe that this bias leads biomedical researchers to focus on richly annotated genes instead of those with the strongest molecular data. We advocate for researchers to reduce these biases by pursuing data-driven hypotheses.

## 2 Introduction

After analyzing samples with a high throughput technology, the de facto first step is to perform pathway or network analysis to identify biological processes that are statistically enriched in the data.^1^ Researchers typically form hypotheses for their follow up experiments based on the genes or proteins involved in the enriched processes. Commonly used resources for identifying gene functions and interactions include the Gene Ontology (GO),^2^ Reactome,^3^ Comparative Toxi-cogenomics Database (CTD),^4^ DrugBank,^5^ Protein Data Bank (PDB),^6^ Pubpular,^7^ and NCBI GeneRIF. Since these resources are created by curation of the scientific literature, they typically only contain functional annotations for genes with published experimental data. Although GO includes predicted functional annotations for genes, they are considered of low quality.^8^ Consequently, researchers select those genes or proteins for further validation that have prior experimental evidence, which, in turn, leads to more functional annotations for those genes at the expense of under-studied genes.

We hypothesized that this experimental paradigm has led to a gene-centric disease research bias where hypotheses are confounded by the streetlight effect of looking for “answers where the light is better rather than where the truth is more likely to lie”.^9–11^ To test this hypothesis, we examined the annotation inequality for the human genome across a number of biomedical databases using gini coefficient, which is a measure of inequality such that high coefficient value indicates higher inequality.^12^

## 3 Results

### 3.1 Gene annotation inequality persists across databases

Despite the tremendous growth of GO from 20,826 annotations in 2004 for 7,524 human genes to 122,926 annotations for 16,173 genes in 2017, annotation inequality in GO has increased from a gini coefficient of 0.34 in 2004 to 0.50 in 2017. The growth in inequality over time validates that genes with existing annotations continue to receive even more annotations. Pathway databases, including Reactome (gini=0.33)^3^ and the CTD (gini=0.47),^4^ have a similarly high level of inequality. Indeed, every gene annotation resource we examined displayed a similarly high level of annotation inequality, including: CTD chemical-gene associations (gini=0.63);^4^ PDB 3D protein structures (gini=0.68);^6^ DrugBank drug-gene associations (gini=0.70);^5^ GeneRIF gene publication annotations (gini=0.79); and Pubpular disease-gene publication associations (gini=0.82).^7, 13^ We calculated global gene annotation gini coefficient of 0.63 when considering the number of annotations pooled across all these databases. When comparing annotation inequality in gene resources to income inequality in the world, we observed that the inequality index for many of the gene resources is higher than any nation in the Organisation for Economic Co-operation and Development (OECD).^14^

### 3.2 Annotation inequality bias affects biomedical research

Next, we explored whether disease research may be affected by the inequality in gene annotation databases. General concern that most published findings are false,^15^ many results are inflated,^16^ and research funding is being wasted^17, 18^ has led to searches for solutions that will yield reproducible and clinically relevant findings. Using a multi-cohort analysis framework,^19, 20^ we have repeatedly demonstrated that it can identify novel disease-gene relationships that lay outside the “halo of the streetlight”, and which have diagnostic, prognostic, and therapeutic utility across diverse diseases including cancer,^21–23^ organ transplantation,^19^ infectious diseases,^24–27^ and autoimmunity.^28^

In our manually curated meta-analyses of 104 distinct human conditions, we have integrated transcriptome data from over 41,000 patients and 619 studies to calculate an effect size for disease-gene associations.^20^ Our meta-analyses covered diverse classes of human conditions, such as cancer, autoimmune disease, viral infection, neurodegenerative and psychiatric disorders, pregnancy, and obesity. For these conditions, we extracted all disease gene associations with at least ten publications.^7, 13^ Published disease-gene associations exhibited no significant correlation with differential gene expression false discovery rate (FDR) rank [Figure 2A, Spearman’s correlation= −0.005, p = 0.716]. Overall, only 19.5% of published disease-gene associations were identified in gene expression meta-analyses at a FDR of 5% [Figure S1a]. This result is consistent with previous publications that have successfully replicated between 11% – 25% of research studies.^29, 30^

**Figure 1:**
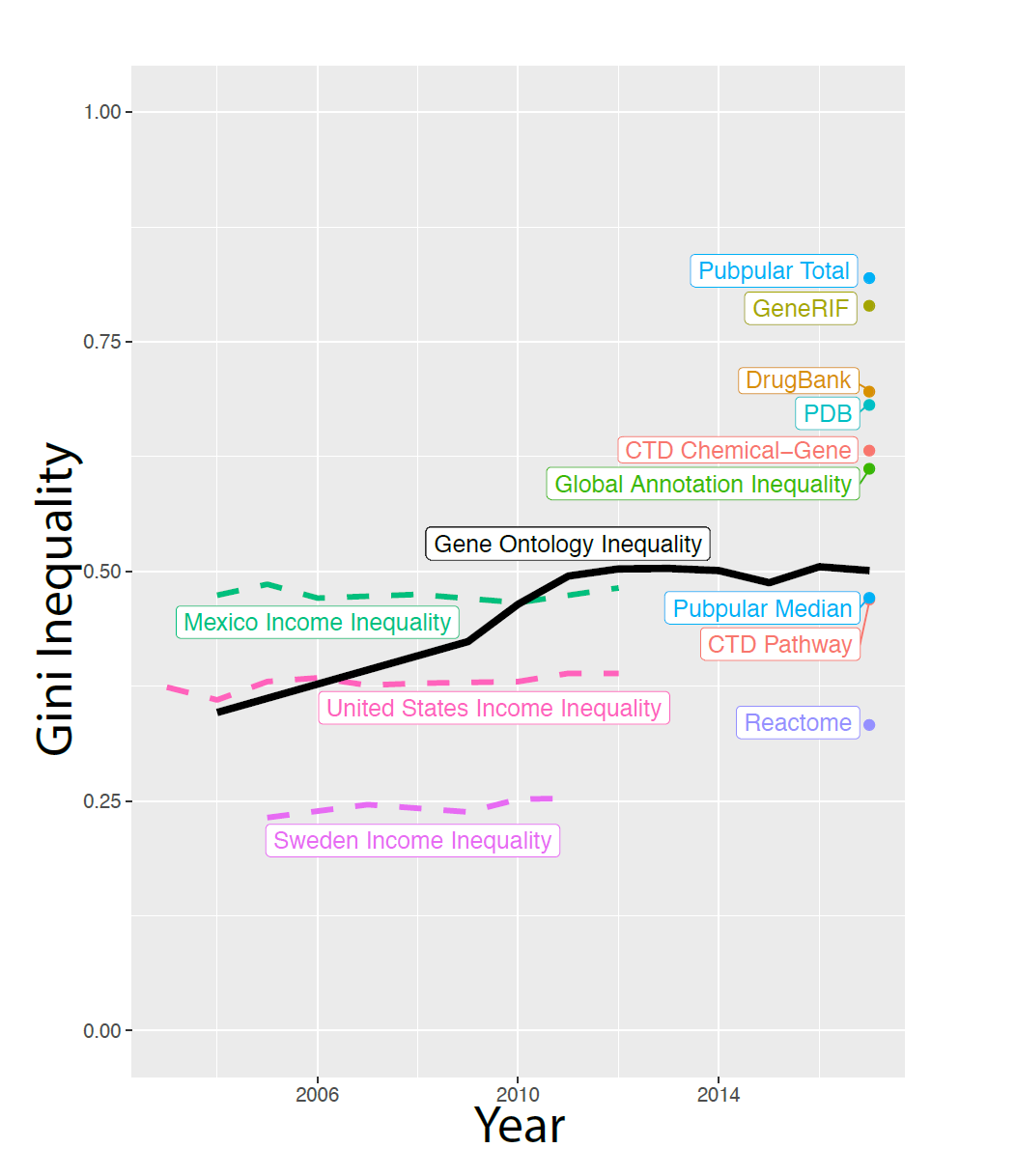
Inequality in gene annotations. We measured the gini coefficient across a variety of gene annotation resources. For comparison, we also displayed gini coefficients for income inequality across a sample of OECD nations.

**Figure 2:**
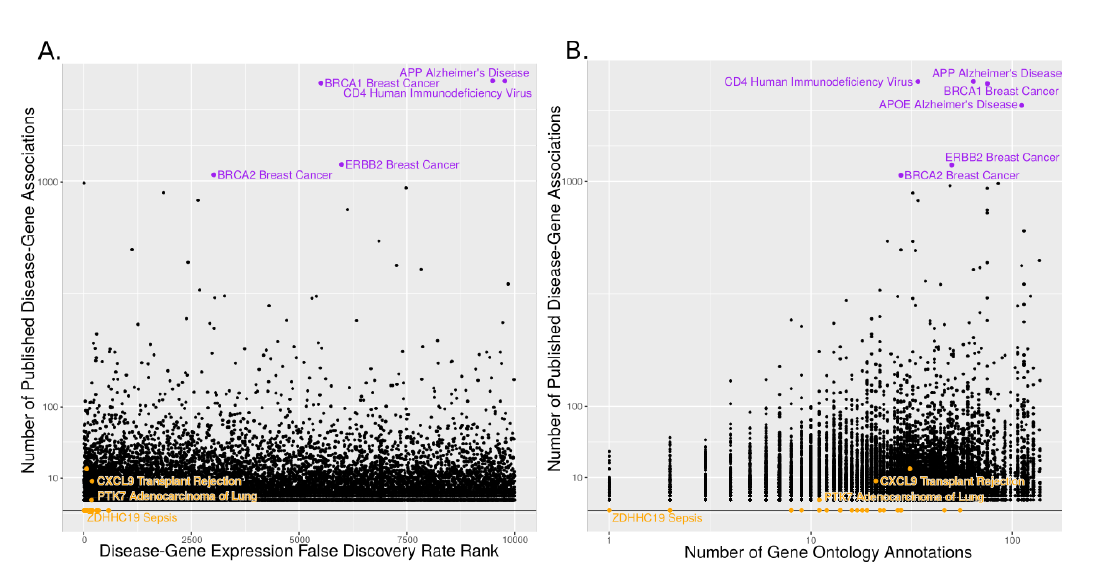
Published Disease-Gene Associations Not Reflected in Molecular Data. (A) The number of publications for every disease-gene pair was not significantly correlated with the gene expression meta-analysis effect size FDR rank [Spearman’s correlation = −0.005, p = 0.716]. (B) The number of publications for every disease-gene pair correlated with the number of non-inferred from electronic annotation (non-IEA) Gene Ontology annotations [Spearman’s correlation = 0.100, p=8.7e-13]. Orange points represent disease-gene associations published in our prior meta-analyses.^19, 23, 24^ Purple points have at least 1000 publications. See also Figure S1.

To observe whether this phenomenon was specific to gene expression, we extracted genome wide significant single nucleotide polymorphisms (SNPs) from the GWAS catalog.^31^ We observed a nominally significant correlation between the number of publications and SNP pvalues, indicating moderate concordance between genetic mutations and disease-gene publications [Figure S1b, Spearman’s correlation = −0.127, p = 0.015].

Based on these results, we hypothesized that the lack of correlation with molecular evidence may have been an artifact of research bias towards well-characterized genes. Therefore, we examined correspondence between publications about a disease-gene pair and existing knowledge about that gene as indicated by the number of GO annotations. Indeed, the number of GO annotations for a gene of interest was significantly correlated with the published disease-gene associations [Figure 2B, Spearman’s correlation = 0.100, p=8.7e-13], but not with gene expression effect size FDR rank in disease [Figure S1c, Spearman’s correlation = −0.010, p = 0.136].^2^ Many of the highly published disease-gene associations may have been studied for reasons that would not be directly reflected in gene expression analysis, including BRCA1 in breast cancer and CD4 in human immunodeficiency virus. The more troubling bias occurs when associations with strong molecular evidence have no publication record. Disease-gene associations we have reported in our published meta-analyses were typically novel findings with few Gene Ontology annotations, despite having extremely low false discovery rates and high effect sizes^19, 21, 24^ [orange points in Figure 2]. We observed similar patterns when we performed the same analysis on similar publication and GWAS data from HuGE Navigator^32, 33^[Figures S1d, S1e, S1f].

## 4 Discussion

Collectively, our results provide evidence of a strong research bias in literature that focuses on well annotated genes instead of the genes with the most significant disease relationship in terms of both gene expression and genetic variation. While focusing research on the best characterized genes may be natural because it is easy to formulate a mechanistic hypothesis of the gene’s function in disease, we propose that omics-era researchers should instead allow data to drive their hypotheses. Our prior work shows that expanding research outside of the streetlight of well characterized genes identifies novel disease-gene relationships, leads to successful repurposings of drugs, and provides clinically actionable diagnostics.^19, 21–27, 34^ To enable researchers to pursue data-driven hypotheses, we have made our gene expression meta-analysis data publicly available at [http://metasignature.stanford.edu] where it may be explored based on either diseases or genes of interest. By focusing on genes with the strongest molecular evidence instead of the most annotations, researchers will break the self-perpetuating annotation inequality cycle that results in research bias.

## 5 Materials and Methods

### 5.1 Gini coefficient calculation

We calculated the gini coefficients using the R package ineq.^35^ We included all human genes with at least one annotation in the gini calculations. We used the Entrez Gene list downloaded in February 2017 of 20,698 current, protein-coding, human genes as our source of human genes.

We calculated the number of annotations for each human gene in the Gene Ontology.^2^ We only considered the biological process and molecular function categories and excluded terms with evidence codes IEA and ND. Duplicate annotations that only differ in evidence codes were counted once. We calculated the number of annotations in the January 2004 release and annually for the January 2009-2017 releases.

We manually downloaded gene-publication data in August 2016 from Pubpular for 102 of the diseases in our gene expression database.^7, 13^ “Pubpular Total” refers to the inequality of gene-publication data across all diseases. “Pubpular Median” refers to the median inequality of gene-publication for each disease.

We downloaded Reactome pathway data from the complete database release 59.^3^ We downloaded data in MySQL format and parsed pathways into UniProt identifiers using custom scripts. We converted UniProt identifiers to gene names using the UniProt identifier conversion tool.^36^ We calculated the number of pathways including each gene name.

We downloaded the CTD^4^ data in February 2017, with the chemical-gene associations and the gene-pathway associations. We calculated the number of chemical-gene and gene-pathway associations for each gene name.

We downloaded GeneRIFs from the NCBI in February 2017. We included all human GeneRIFS (Tax ID: 9606). We calculated the number of GeneRIFs for each gene.

We downloaded the gene names associated with protein structures from the Protein Data Bank^6^ in February 2017 and calculated the number of structures per gene name.

We downloaded the DrugBank^5^ database version 5.0.5 and identified all drugs with known activities on human genes. We calculated the number of drugs targeting each gene.

We downloaded OECD^14^ nation income inequality gini coefficients from the July 2016 data release at http://www.oecd.org/social/income-distribution-database. htm.

The code and data we used to run this analysis is available at http://khatrilab.stanford.edu/researchbias.

### 5.2 Gene expression data collection and meta-analysis

Gene expression meta-analysis data was compiled from the MetaSignature database.^20^ MetaSignature includes data from manual meta-analysis of over 41,000 samples, 619 studies, and 104 diseases. Briefly, relevant data were downloaded from Gene Expression Omnibus and Array-Express.^37, 38^ Cases and controls were manually labeled for each disease and meta-analysis was performed using the MetaIntegrator package.^20^ We used the Hedges’ *g* summary effect size, standard error, and false discovery rate which the MetaIntegrator package calculates for every gene.

### 5.3 Data collection for disease-gene publications and SNP data

We downloaded the number of publications for each disease-gene relationship from PubPular and HuGE Navigator in August 2016 for as many of the 104 disease in MetaSignature as were present in the databases (102 in PubPular and 81 in HuGE).^7, 13, 32^ PubPular gave the top 261 gene associations, and HuGE gave all known associations. For all correlations, we only considered disease-gene associations with at least 10 publications to limit false positive associations.

We downloaded disease-SNP relationships, including gene mappings, odds ratios, and p-values, from the GWAS Catalog and HuGE Navigator for 61 and 54, respectively, of the 103 diseases in MetaSignature.^31, 33^ From Gene Ontology, we calculated the counts of non-Inferred from Electronic Annotation annotations for all the genes in the MetaSignature database.^2^ The Spearman rank correlation was used for all correlations.

Our plots show the top 10,000 gene associations for each disease by effect size FDR rank. Correlation calculations do not include a similar limit.

## 6 Acknowledgements

We thank Paul J. Utz for feedback about the manuscript and figures and Alex Schrenchuk for computer support. WAH is funded by the National Science Foundation Graduate Research Fellowship under Grant No. DGE-114747. PK is funded by the the Bill and Melinda Gates Foundation, and NIAID grants 1U19AI109662, U19AI057229 and U54I117925.

## 7 Author Contributions

Conceptualization, W.A.H., A.T., and P.K; Methodology, W.A.H., A.T., and P.K; Software, W.A.H. and A.T.; Investigation, W.A.H. and A.T.; Data Curation, W.A.H. and A.T.; Writing-Original Draft, W.A.H. and P.K.; Writing-Reviewing and Editing, W.A.H., A.T., and P.K.; Visualization, W.H.; Funding Acquisition, P.K.

## 8 Supplementary Materials

**Figure S1:**
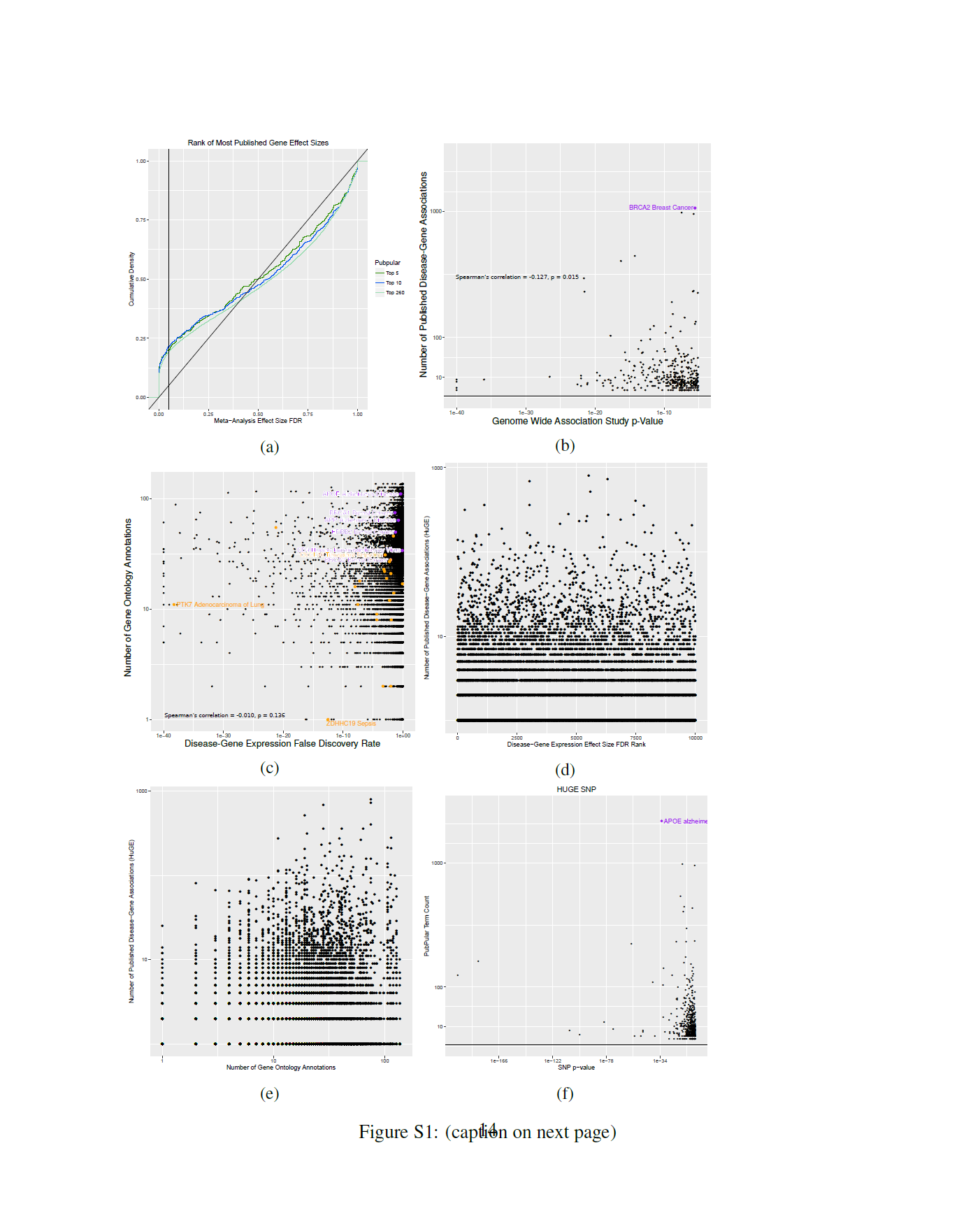
Related to Figure 2. (a) Only 20% of published disease-gene associations have gene effect size FDR of less than 5%. Cumulative distributions of the top 5, 10, and 250 diseasegene associations for each disease from PubPular database. Vertical line at a gene expression meta-analysis effect size FDR of 5%. (b) The number of publications for every disease-gene pair is nominally significantly correlated with published results from SNP GWAS from the GWAS catalog [Spearman’s correlation = −0.127, p = 0.015]. (c) The number of gene ontology annotations for every gene is not correlated with the gene expression meta-analysis effect size false discovery rate (FDR) rank [Spearman’s correlation = −0.010, p = 0.156]. (d) The number of publications for every disease-gene pair vs. the gene expression meta-analysis effect size FDR rank based on the HuGE Navigator data [Spearman’s correlation = 0.032, p = 0.169]. (e) The number of publications for every disease-gene pair vs. the number of non-inferred from electronic annotation (non-IEA) Gene Ontology annotations based on the HuGE Navigator data [Spearman’s correlation = 0.211, p = 2.2e-16]. (f) The number of publications for every disease-gene pair vs. SNP GWAS p-value based on the HuGE Navigator data [Spearman’s correlation = −0.127, p = 0.015].

## References and Notes

1 Khatri, P., Sirota, M. & Butte, A. J. Ten years of pathway analysis: current approaches and outstanding challenges. PLoS computational biology 8, e1002375 (2012). URL http://journals.plos.org/ploscompbiol/article?id=10.1371/journal.pcbi.1002375\#pcbi-1002375-g003.

2 Ashburner, M. et al. Gene ontology: tool for the unification of biology. The Gene Ontology Consortium. Nature genetics 25, 25–9 (2000). URL http://dx.doi.org/10.1038/75556.

3 Croft, D. et al. The Reactome pathway knowledgebase. Nucleic Acids research 42, D472–7 (2014). URL http://nar.oxfordjournals.org/content/42/D1/D472.abstract.

4 Davis, A. P. et al. The Comparative Toxicogenomics Database’s 10th year anniversary: update 2015. Nucleic Acids research 43, D914–20 (2015). URL http://nar.oxfordjournals.org/content/43/D1/D914.short.

5 Wishart, D. S. et al. DrugBank: a comprehensive resource for in silico drug discovery and exploration. Nucleic Acids research 34, D668–72 (2006). URL http://www.pubmedcentral.nih.gov/articlerender.fcgi?artid=1347430\&tool=pmcentrez\&rendertype=abstract.

6 Berman, H. M. et al. The Protein Data Bank. Nucleic Acids research 28, 235–42 (2000). URL http://www.ncbi.nlm.nih.gov/pubmed/10592235 http://www.pubmedcentral.nih.gov/articlerender.fcgi?artid=PMC102472.

7 Maggie Lam. PubPular: Identifying the focus of biomedical research. URL https://pubpular.shinyapps.io/PubPular/.

8 Yon Rhee, S., Wood, V., Dolinski, K. & Draghici, S. Use and misuse of the gene ontology annotations. Nature Reviews Genetics 9, 509–515 (2008). URL http://www.nature.com/doifinder/10.1038/nrg2363.

9 Freedman, D. H. Why Scientific Studies Are So Often Wrong: The Streetlight Effect. Dis188 cover Magazine 1 (2010).

10 Battaglia, M. & Atkinson, M. A. The streetlight effect in type 1 diabetes. Diabetes 64, 1081–90 (2015). URL http://www.ncbi.nlm.nih.gov/pubmed/25805758 www.pubmedcentral.nih.gov/articlerender.fcgi?artid=PMC4375074.

11 Bulgheresi, S. Bacterial cell biology outside the streetlight. Environmental Microbiology 18, 2305–2318 (2016). URL http://doi.wiley.com/10.1111/1462-2920.13406.

12 Gini, C. & C. Variabilitàe mutabilità. Reprinted in Memorie di metodologica statistica (Ed. Pizetti E, Salvemini, T). Rome: Libreria Eredi Virgilio Veschi (1912).

13 Lam, M. P. Y. et al. Data-Driven Approach To Determine Popular Proteins for Targeted Proteomics Translation of Six Organ Systems. Journal of proteome research Web (2016). URL http://www.ncbi.nlm.nih.gov/pubmed/27356587.

14 OECD Income Distribution Database (2016). URL http://www.oecd.org/social/income-distribution-database.htm.

15 Ioannidis, J. P. A. Why most published research findings are false. PLoS medicine 2, e124 (2005). URL http://journals.plos.org/plosmedicine/article?id=10.1371/journal.pmed.0020124.

16 Ioannidis, J. P. A. Why Most Discovered True Associations Are Inflated. Epidemiology 19, 640–648 (2008). URL http://content.wkhealth.com/linkback/openurl?sid=WKPTLP:landingpage\&an=00001648-200809000-00002.

17 Macleod, M. R. et al. Biomedical research: increasing value, reducing waste (2014).

18 Collins, F. S. & Tabak, L. A. Policy: NIH plans to enhance reproducibility. Nature 505, 612–613 (2014). URL http://www.nature.com/doifinder/10.1038/505612a.

19 Khatri, P. et al. A common rejection module (CRM) for acute rejection across multiple organs identifies novel therapeutics for organ transplantation. The Journal of experimental medicine 210, 2205–21 (2013). URL http://jem.rupress.org/content/210/11/2205.full.

20 Haynes, W. A. et al. Empowering Multi-Cohort Gene Expression Analysis to Increase Reproducibility. bioRxiv Web (2016). URL http://biorxiv.org/content/early/2016/08/25/071514.

21 Mazur, P. K. et al. SMYD3 links lysine methylation of MAP3K2 to Ras-driven cancer. Nature advance on (2014). URL www.nature.com/articles/nature13320.

22 Mazur, P. K. et al. Combined inhibition of BET family proteins and histone deacetylases as a potential epigenetics-based therapy for pancreatic ductal adenocarcinoma. Nature Medicine 21, 1163–1171 (2015). URL http://www.nature.com/doifinder/10.1038/ nm.3952.

23 Chen, R. et al. A meta-analysis of lung cancer gene expression identifies PTK7 as a survival gene in lung adenocarcinoma. Cancer Research 74, 2892–2902 (2014). URL http://www.ncbi.nlm.nih.gov/pubmed/24654231.

24 Sweeney, T. E., Shidham, A., Wong, H. R. & Khatri, P. A comprehensive time-course-based multicohort analysis of sepsis and sterile inflammation reveals a robust diagnostic gene set. Science Translational Medicine 7, 287ra71 (2015). URL http://stm.sciencemag.org/content/7/287/287ra71.abstract.

25 Andres-Terre, M. et al. Integrated, Multi-cohort Analysis Identifies Conserved Transcriptional Signatures across Multiple Respiratory Viruses. Immunity 43, 1199–1211 (2015). URL http://www.cell.com/article/S1074761315004550/fulltext.

26 Sweeney, T. E., Braviak, L., Tato, C. M. & Khatri, P. Genome-wide expression for diagnosis of pulmonary tuberculosis: a multicohort analysis. The Lancet Respiratory Medicine 4, 213–224 (2016).

27 Sweeney, T. E.,Wong, H. R. & Khatri, P. Robust classification of bacterial and viral infections via integrated host gene expression diagnostics. Science translational medicine 8, 346ra91 (2016). URL http://www.ncbi.nlm.nih.gov/pubmed/27384347.

28 Lofgren, S. et al. Integrated, multicohort analysis of systemic sclerosis identifies robust transcriptional signature of disease severity. JCI Insight 1 (2016). URL https://insight.jci.org/articles/view/89073.

29 Prinz, F., Schlange, T. & Asadullah, K. Believe it or not: how much can we rely on published data on potential drug targets? Nature Reviews Drug Discovery 10, 712–712 (2011). URL http://www.nature.com/doifinder/10.1038/nrd3439-c1.

30 Begley, C. G. & Ellis, L. M. Drug development: Raise standards for preclinical cancer research. Nature 483, 531–3 (2012). URL http://www.nature.com/nature/journal/v483/n7391/full/483531a.html\#t1.

31 Welter, D. et al. The NHGRI GWAS Catalog, a curated resource of SNP-trait associations. Nucleic Acids research 42, D1001–6 (2014). URL http://www.ncbi.nlm.nih.gov/pubmed/24316577 http://www.pubmedcentral.nih.gov/articlerender.fcgi?artid=PMC3965119.

32 Yu, W., Clyne, M., Khoury, M. J. & Gwinn, M. Phenopedia and Genopedia: disease-centered and gene-centered views of the evolving knowledge of human genetic associations. Bioinformatics 26, 145–146 (2010). URL http://bioinformatics.oxfordjournals.org/cgi/doi/10.1093/bioinformatics/btp618.

33 Yu, W. et al. GWAS Integrator: a bioinformatics tool to explore human genetic associations reported in published genome-wide association studies. European Journal of Human Genetics 19, 1095–1099 (2011). URL http://www.nature.com/doifinder/10.1038/ejhg.2011.91.

34 Li, M. D., Burns, T. C., Morgan, A. A. & Khatri, P. Integrated multi-cohort transcriptional meta-analysis of neurodegenerative diseases. Acta neuropathologica communications 2, 93 (2014). URL http://www.pubmedcentral.nih.gov/articlerender.fcgi?artid=4167139\&tool=pmcentrez\&rendertype=abstract

35 Zeileis, A. ineq: Measuring Inequality, Concentration, and Poverty (2014). URL https://cran.r-project.org/package=ineq.

36 UniProt: the universal protein knowledgebase. Nucleic Acids Research 45, D158–D169 (2017). URL https://academic.oup.com/nar/article-lookup/doi/10.1093/nar/gkw1099.

37 Brazma, A. et al. ArrayExpress-a public repository for microarray gene ex270 pression data at the EBI. Nucleic Acids Research 31, 68–71 (2003). URL http://www.pubmedcentral.nih.gov/articlerender.fcgi?artid=165538\&tool=pmcentrez\&rendertype=abstract.

38 Edgar, R. Gene Expression Omnibus: NCBI gene expression and hybridization array data repository. Nucleic Acids Research 30, 207–210 (2002). URL http://nar.oxfordjournals.org/content/30/1/207.short.

